# FlashFry: a fast and flexible tool for large-scale CRISPR target design

**DOI:** 10.1101/189068

**Authors:** Aaron McKenna, Jay Shendure

**Affiliations:** Genome Sciences, University of Washington, Seattle, WA, USA; Howard Hughes Medical Institute, Seattle WA, USA

## Abstract

FlashFry is a fast and flexible command-line tool for characterizing large numbers of CRISPR target sequences. While several CRISPR web application exist, genome-wide knockout studies, noncoding deletion scans, and other large-scale studies or methods development projects require a simple and lightweight framework that can quickly discover and score thousands of candidates guides targeting an arbitrary DNA sequence. With FlashFry, users can specify an unconstrained number of mismatches to putative off-targets, richly annotate discovered sites, and tag potential guides with commonly used on target and off-target scoring metrics. FlashFry runs at speeds comparable to widely used genome-wide sequence aligners, and output is provided as an easy-to-manipulate text file.

**Availability:** FlashFry is written in Scala and bundled as a stand-alone Jar file, easily run on any system with an installed Java virtual machine (JVM). The tool is freely licensed under version 3 of the GPL, and code, documentation, and tutorials are available on the GitHub page: http://aaronmck.github.io/FlashFry/

## 1 Introduction

The CRISPR prokaryotic immune system has transformed genome engineering. As typically used, CRISPR proteins are directed to create double-stranded DNA breaks at location(s) in a genome matching a specified guide sequence (Wright *et al.* (2016)). These double-stranded breaks are commonly repaired by a non-homologous end joining (NHEJ) pathway, which can leave small insertions or deletions (indels) at the genomic target site. The site-specific introduction of such indels can be used to perturb endogenous gene function (Wang *et al*. (2017)), encode information (McKenna *et al.* (2016)), or characterize the function of genomic sequence (Gasperini *et al*. (2017); Chen *et al*. (2015); Liu *et al*. (2017)).

Although CRISPR editing is highly specific (Doench *et al*. (2014)), not all guides function with the same efficiency or specificity. For instance, double-stranded breaks can occur at genomic locations (‘targets’) that are an imperfect match to the supplied guide sequence (termed ‘off-targets’). To reduce the chance of such unintended genome editing, guide sequences can be chosen that contain less overlap with all possible alternate targets in the genome. The importance of specific differences in the guide sequence, the genomic location and chromatin environment of the target, and the method of guide delivery all have effects on the distribution and rate of off-target cutting (Haeussler *et al*. (2016)).

To help users choose both specific and active guide sequences, the community has created a large number of CRISPR design tools, most of which are made available as web applications (Labun *et al*. (2016); Haeussler *et al*. (2016)). These web tools are convenient for researchers screening a small set of guides, or scanning a single genomic locus like an exon. Unfortunately, these tools require batched queries for large sets, which makes it more challenging to scan loci or whole genomes for guides. Additionally, some guide screening tools rely upon genome-wide alignment tools to generate putative off-target lists for each guide. For practical reasons, these aligners are generally designed to quickly discover only the most similar sequences with a limited number of mismatches in comparison to the guide (typically *k* ≤ 3), whereas experimental efforts have shown activity at off-target sequences containing upwards of six mismatches to the guide (Tsai *et al*. (2014)). Some of these tools also miss a subset of potential off-targets altogether, regardless of the mismatch distance (Doench *et al*. (2016)). To address these issues, and to meet our needs for high-throughput guide selection from arbitrary genomic regions, we’ve created FlashFry, a command-line tool for discovery and characterization of CRISPR guide sequences.

## 2 Materials and methods

### 2.1 Database creation

FlashFry generates a block-compressed binary database of all potential CRISPR targets for a given reference sequence. This database is CRISPR enzyme specific, and can be generated in a few hours on a standard computer (supplemental table 1). In this database, target sequences that contain the CRISPR enzyme’s protospacer adjacent motif (PAM) are encoded with their positions and counts into a hierarchy of sorted prefix-bins (figure 1A). This prefix length can be specified at runtime, with larger bins being automatically sub-indexed to reduce lookup times. Within each bin, target sequences and the number of occurrences in the genome are stored as a 64 bit-encoded value. This is followed by additional binary encoded values for each target’s position within the genome.

**Fig. 1.**
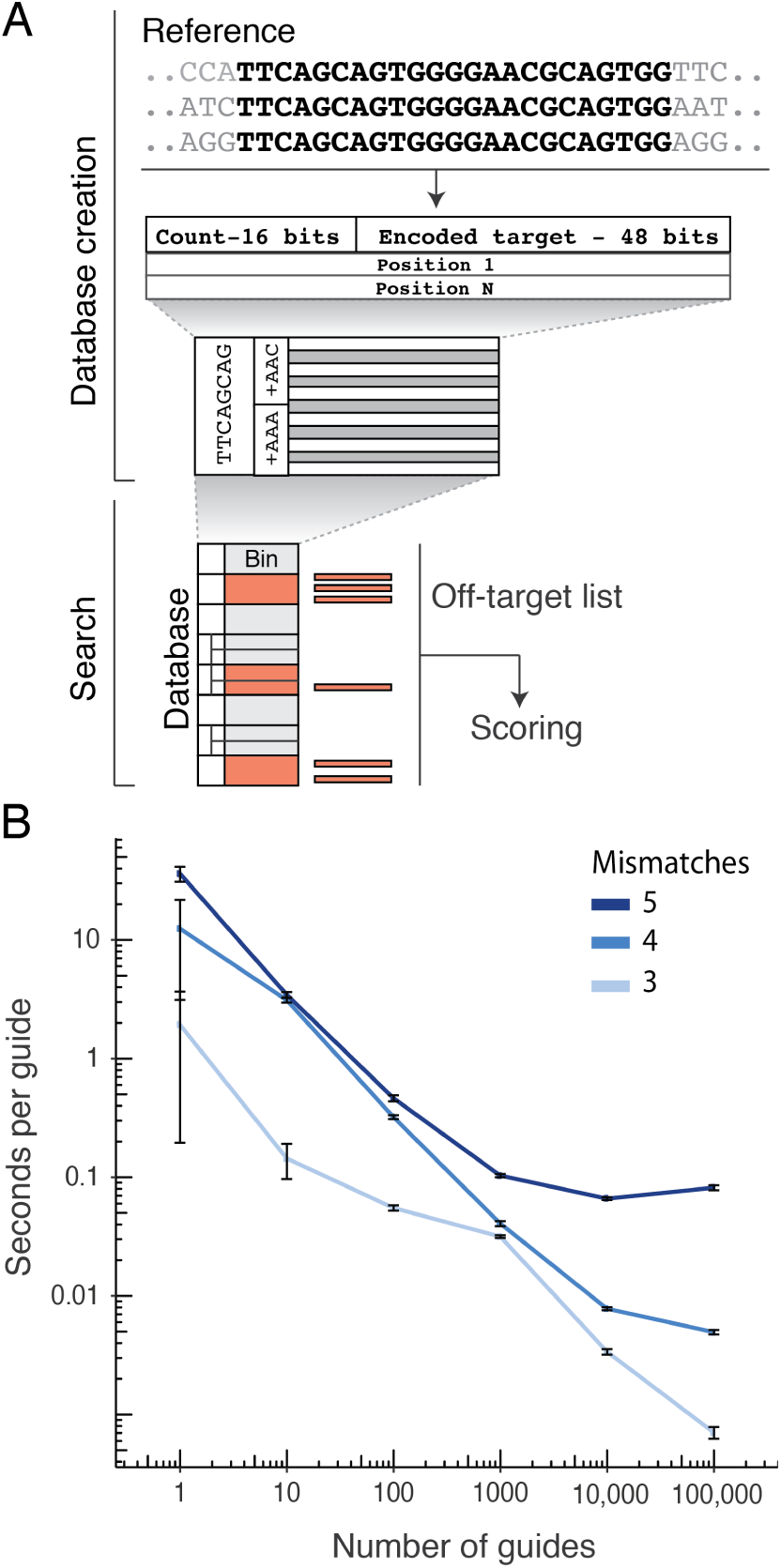
**(A)** FlashFry schematic. The genome of interest is scanned for targets that match the PAM of the specified CRISPR enzyme, which are then aggregated by their prefix and bit-encoded into a compressed, hierarchical database of genomic targets. This database can then be searched for off-targets by comparing the guide prefix against the contents of bins within *k* mismatches. **(B)** Median runtime and median absolute deviation (MAD) of batched random guide sequences (twenty replicates) with an increasing number of allowed mismatches to targets.

### 2.2 Search

Given the inherent inefficiencies of high-mismatch searches, FlashFry uses a filtering approach to find candidate off-targets. It does so by precomputing a traversal over prefix bins with less than *k* mismatches to each guide in the candidate set using a prefix tree (such filtering approaches are reviewed in Navarro (2001)). When a large number of bins are to be searched, which is common with large guide screens or with a high *k* mismatch threshold, FlashFry will instead search the full database to avoid the cost of disk seeks. To further reduce search times, target sequences and their occurrences are stored as a bit-encoded value, allowing bit-parallelism comparisons when determining mismatches (Navarro (2001)), supplementary figure 1). FlashFry is compatible with target sequences up to 24 bases in length, although it could be expanded to longer target sequences as the bins encode their prefix. Lastly, off-target discovery is halted for candidate guides that have exceeded a user defined number of off-target hits, saving compute time by eliminating poor candidates early from the putative guide pool. Per-guide search times decrease with the number of guides and the allowed number of mismatches (figure 1B), and compare favorably to similar CRISPR command line tools or FM-index based tools (supplementary figure 2).

### 2.3 Guide characterization and scoring

The goal for most users is to pick some subset of highly active and specific guide sequences from a full list of candidate targets within a region of interest. Therefore, FlashFry reports many commonly used scoring approaches for both on-target efficiency as well as off-target performance, including Cutting-frequency determination (CFD)(Doench *et al*. (2016)) the Hsu *et. al*. off-target scoring scheme (Hsu *et al*. (2013)), and both the Moreno-Mateos and Vejnar *et. al*. and the Doench *et al.* 2014 on-target metrics (Moreno-Mateos *et al*. (2015); Doench *et al*. (2014)). We have also included a set of basic design criteria filters, including high and low GC content, warnings for poly T tracts (which halt Pol. III transcription), and targets that have reciprocal off-targets within the region of interest (potentially leading to deletions of the intervening sequence). Lastly, regions can be annotated with information from external BED files, which may be useful for highlighting repetitive sequences or putative regulatory regions. To demonstrate the utility of FlashFry for creating CRISPR libraries, we scored 254,848 candidate SpCas9 target sequences (allowing both NGG and NAG PAMs) within 1 megabase of the human *MYC* gene (supplementary figure 3). The results were scored with two on-target and two off-target metrics from the literature, and intersected with a list of known repetitive elements. The aggregate table could then be used to design a perturbation screen; for instance, 8,038 candidate guide sequences had no other exact occurrence within the human genome, had a Hsu et. al. score above 70, and did not overlap a repetitive element annotation.

## 3 Discussion

The needs of genome-wide knockout studies, noncoding deletion scans, and other large-scale studies or methods development projects are unfortunately not well-met by the abundant CRISPR web applications. FlashFry, an efficient and flexible toolset, fills this void, and can be used to rapidly discover and characterize tens to hundreds-of-thousands of guides from an arbitrary sequence quickly and with a low memory footprint. For methods developers, we also expose a simple interface for implementing additional scoring schemes, given the sequence context of a guide and its off-target hits. FlashFry has no system dependencies outside of the JVM, and avoids many of the pitfalls and complexity of tools that rely on genome aligners to discover off-target sequences.

## Acknowledgements

We thank the members of the Shendure lab for their feedback, suggestions, and critical reading of the manuscript, especially Molly Gasperini, Vikram Agarwal, and Martin Kircher. This work was funded by grants from the NIH (DP1HG007811 and R01HG006283 to J.S.), the Paul G. Allen Family Foundation (to J.S.). AM was supported by a fellowship from the NIH/NHLBI (T32HL007312). JS is an investigator of the Howard Hughes Medical Institute.

---

**Supplementary Figure 1.**
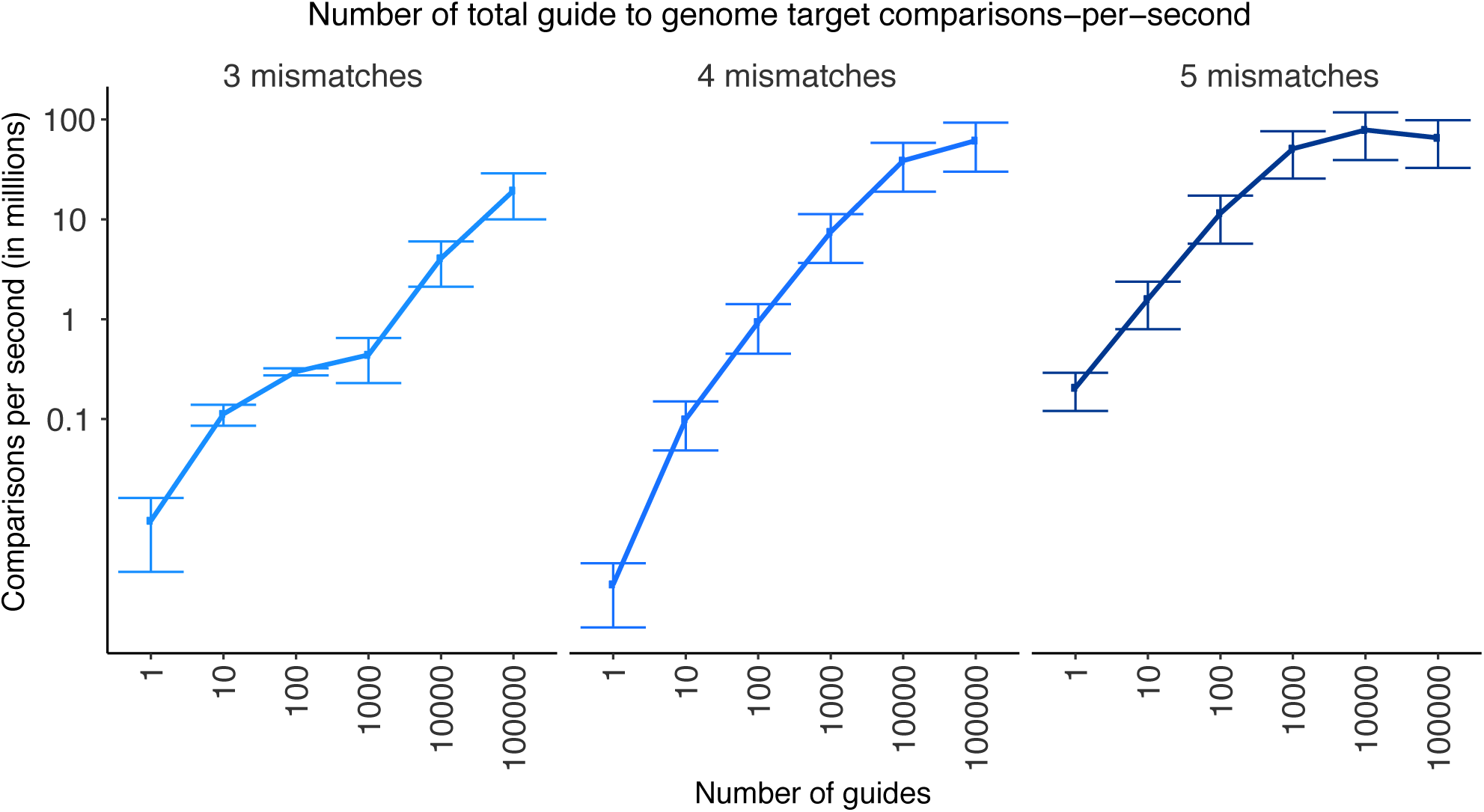
Guide-to-target comparisons per-second. Comparisons per-second for1, 3, and5 allowed mismatches over an increasing number of candidate guides. Smaller numbers of guides and lower mismatch counts achieve lower comparison rates as the initialization and output times are amortized over the whole run.

**Supplementary Figure 2.**
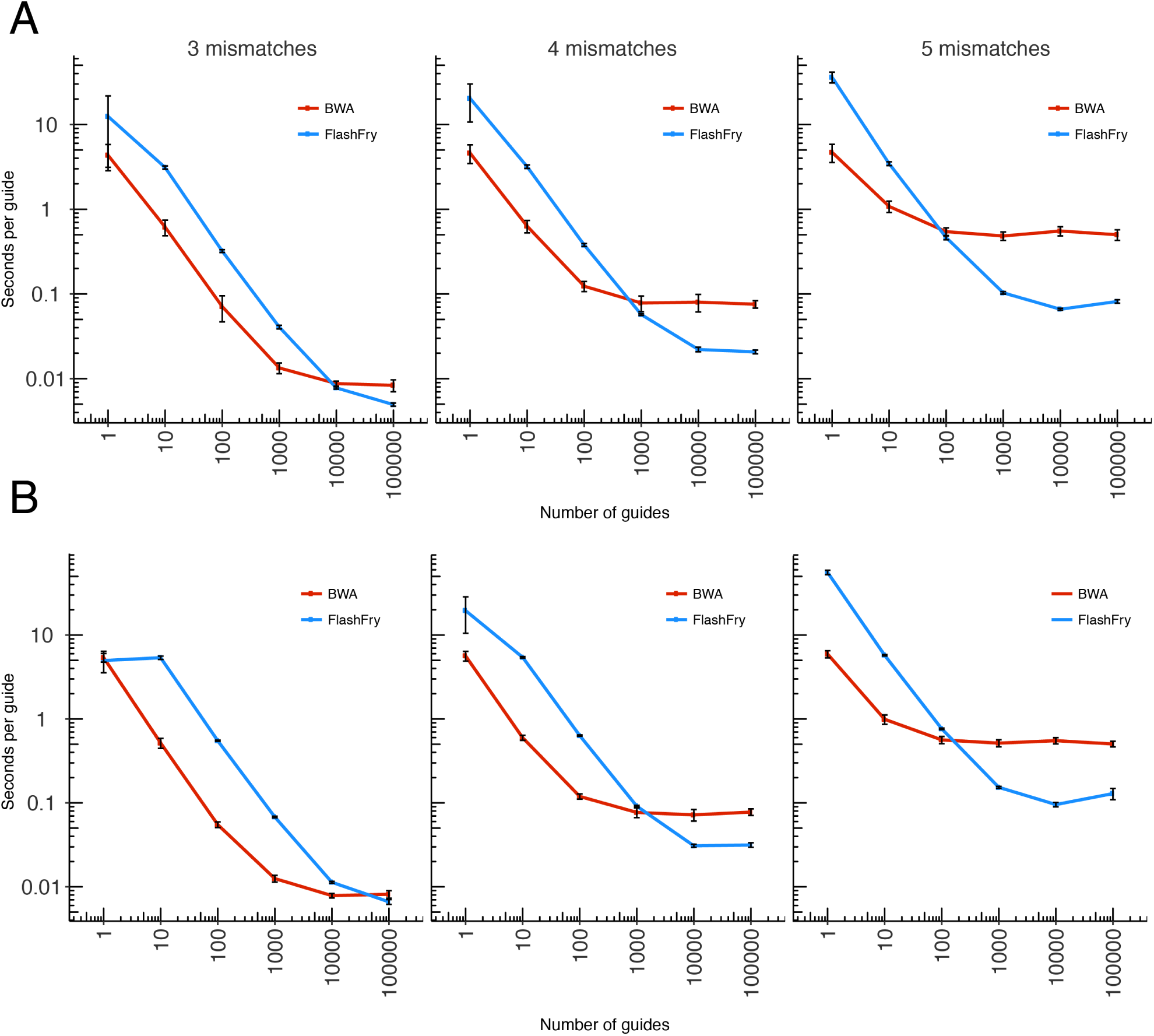
Runtime comparison of FlashFry and BWA. Comparison of runtimes for FlashFry and BWA version 0.7.13-r1126 (Li and Durbin 2009) over an increasing number of guides and permitted mismatches. Twenty-five random CRISPR guide sets were run for each guide-count (x-axis) and permitted mismatch level (2,777,775 potential guides per mismatch level). BWA runtime includes the initial alignment step (aln) and mapping to genomic coordinates (samse), and BWA was run with parameters taken from Haeussler *et al*. (2016): *bwa aln - o 0 - m 20000000 - n mismatches - k mismatches - N - l 20*. Plotted are the median runtime with median absolute deviation (MAD) bars for each set of 25 runs. **(A)** Using the NGG motif for off-target selection, **(B)** using the NRG motif for off-target selection. FlashFry benefits from aggregating all guide-to-genome comparisons in one pass of the database, matching BWA’s performance at hundreds of guides for 5 mismatches, and thousands of guides at 4 mismatches.

**Supplementary Table 1.** Off-target database generation times. A sample of computational times (h:m:s) required to build a FlashFry database for versions of the *Caenorhabditis elegans*, human, mouse, and *Drosophila melanogaster* genomes for common CRISPR enzymes. This analysis was run on a disk-based network area storage (NAS) system.

| Genome | Cas9 (NGG) | Cas9 (NGG or NAG) | CPF1 (TTTN) |
| --- | --- | --- | --- |
| *Caenorhabditis elegans* - 235 | 0:3:21 | 0:6:03 | 0:5:35 |
| Human - hg38 | 3:19:29 | 5:24:55 | 2:50:59 |
| Mouse - mm10 | 2:36:53 | 4:36:03 | 2:11:35 |
| *Drosophila melanogaster*- BDGP6 | 0:6:33 | 0:10:48 | 0:5:44 |

**Supplementary Figure 3.**
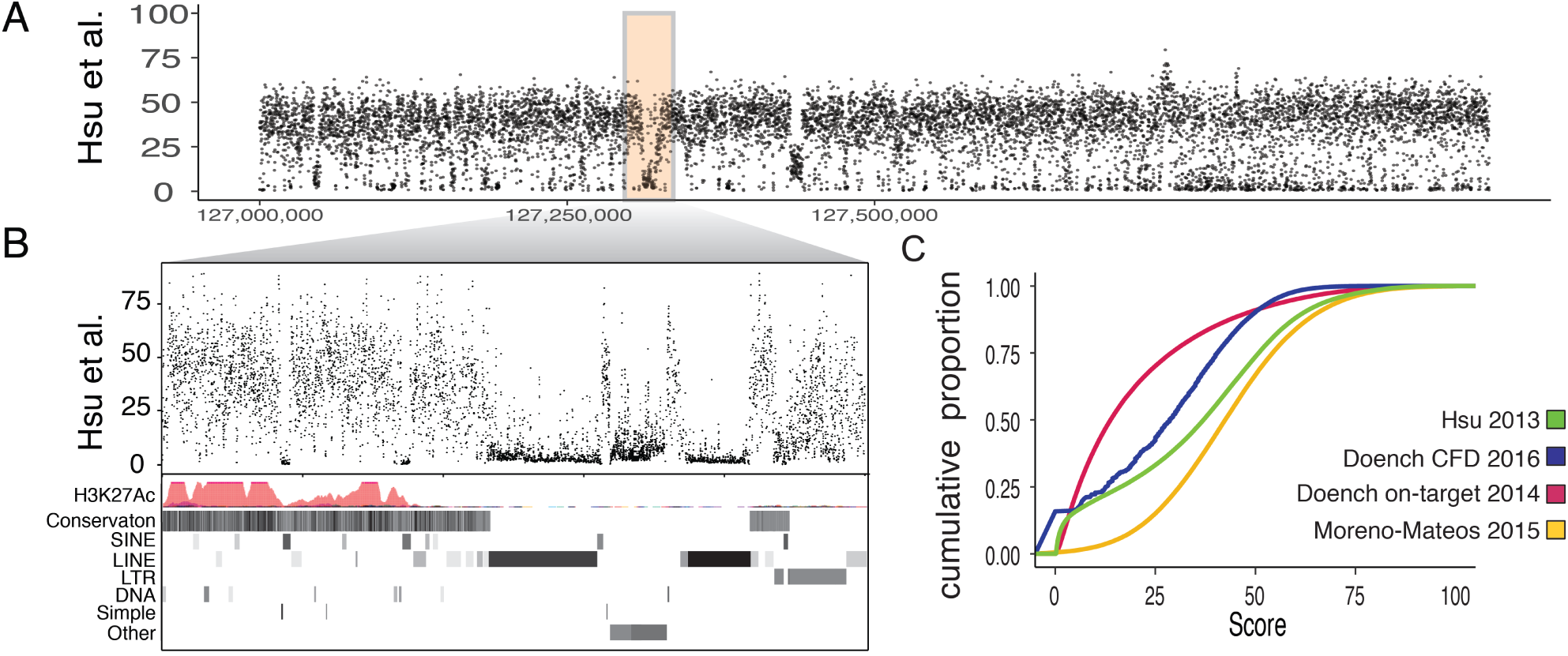
Candidate guide discovery over the human MYC region. **(A)** FlashFry was run for a 1 Mb region flanking the human *MYC* gene (chr8:127,000,000-128,000,000, human reference hg38), generating 254,848 candidate sites, which were scored for both on and off-target activity. Average off-target specificity scores from Hsu *et al*. are are shown, averaged over 100 basepair windows. **(B)** Enlargement of the 25Kb region (chr8:127,300,000-127,325,000) highlighted in **A**. The observed off-target specificity tracks well with known repetitive elements. **(C)** Cumulative density function plot of on and off-target scoring metrics for all targets over the region highlighed in **A**.

**Supplementary Table 2.** Effects of SSD storage on runtime. Off-target discovery times (seconds) for 10,000 random guide sequences on an G2 Amazon EC2 node with an SDD drive for off-target database storage, compared to a distributed node on a local cluster with a disk-based network area storage (NAS). Both jobs were run with identical parameter sets and memory allocations.

| Data set | Amazon EC2 (G2) instance | Local distributed cluster nodes |
| --- | --- | --- |
| 10000 guide (set A) | 351.12 | 776.22 |
| 10000 guide (set B) | 348.53 | 825.57 |
| 10000 guide (set C) | 347.86 | 1397.88 |
| 10000 guide (set D) | 344.26 | 783.44 |
| 10000 guide (set E) | 347.30 | 776.44 |
| mean | 347.81 | 911.91 |
| Standard deviation | 2.46 | 272.47 |

